# From single nuclei to whole genome assemblies

**DOI:** 10.1101/625814

**Authors:** Merce Montoliu-Nerin, Marisol Sánchez-García, Claudia Bergin, Manfred Grabherr, Barbara Ellis, Verena Esther Kutschera, Marcin Kierczak, Hanna Johannesson, Anna Rosling

## Abstract

A large proportion of Earth's biodiversity constitutes organisms that cannot be cultured, have cryptic life-cycles and/or live submerged within their substrates^1–4^. Genomic data are key to unravel both their identity and function^5^. The development of metagenomic methods^6,7^ and the advent of single cell sequencing^8–10^ have revolutionized the study of life and function of cryptic organisms by upending the need for large and pure biological material, and allowing generation of genomic data from complex or limited environmental samples. Genome assemblies from metagenomic data have so far been restricted to organisms with small genomes, such as bacteria^11^, archaea^12^ and certain eukaryotes^13^. On the other hand, single cell technologies have allowed the targeting of unicellular organisms, attaining a better resolution than metagenomics^8,9,14–16^, moreover, it has allowed the genomic study of cells from complex organisms one cell at a time^17,18^. However, single cell genomics are not easily applied to multicellular organisms formed by consortia of diverse taxa, and the generation of specific workflows for sequencing and data analysis is needed to expand genomic research to the entire tree of life, including sponges^19^, lichens^3,20^, intracellular parasites^21,22^, and plant endophytes^23,24^. Among the most important plant endophytes are the obligate mutualistic symbionts, arbuscular mycorrhizal (AM) fungi, that pose an additional challenge with their multinucleate coenocytic mycelia^25^. Here, the development of a novel single nuclei sequencing and assembly workflow is reported. This workflow allows, for the first time, the generation of reference genome assemblies from large scale, unbiased sorted, and sequenced AM fungal nuclei circumventing tedious, and often impossible, culturing efforts. This method opens infinite possibilities for studies of evolution and adaptation in these important plant symbionts and demonstrates that reference genomes can be generated from complex non-model organisms by isolating only a handful of their nuclei.

## Main text

AM fungi is a group of diverse obligate symbionts that have colonized root cells and formed mycelial networks in soil since plants first colonized land^25–27^. Their entire life-cycle is completed underground and they propagate with multinuclear asexual spores^27^ (Figure 1). Genomic research on AM fungi has been hampered by technical challenges involving isolation and culturing, and accordingly, few species have been successfully sequenced. To date, the reference genomes of only few species that can be grown in axenic culture, *i.e., Rhizophagus irregularis*, *R. clarus*, *R. diaphanus*, *R. cerebriforme*, *Gigaspora rosea* and *Diversispora epigaea,* have been published^28–33^.

**Figure 1.**
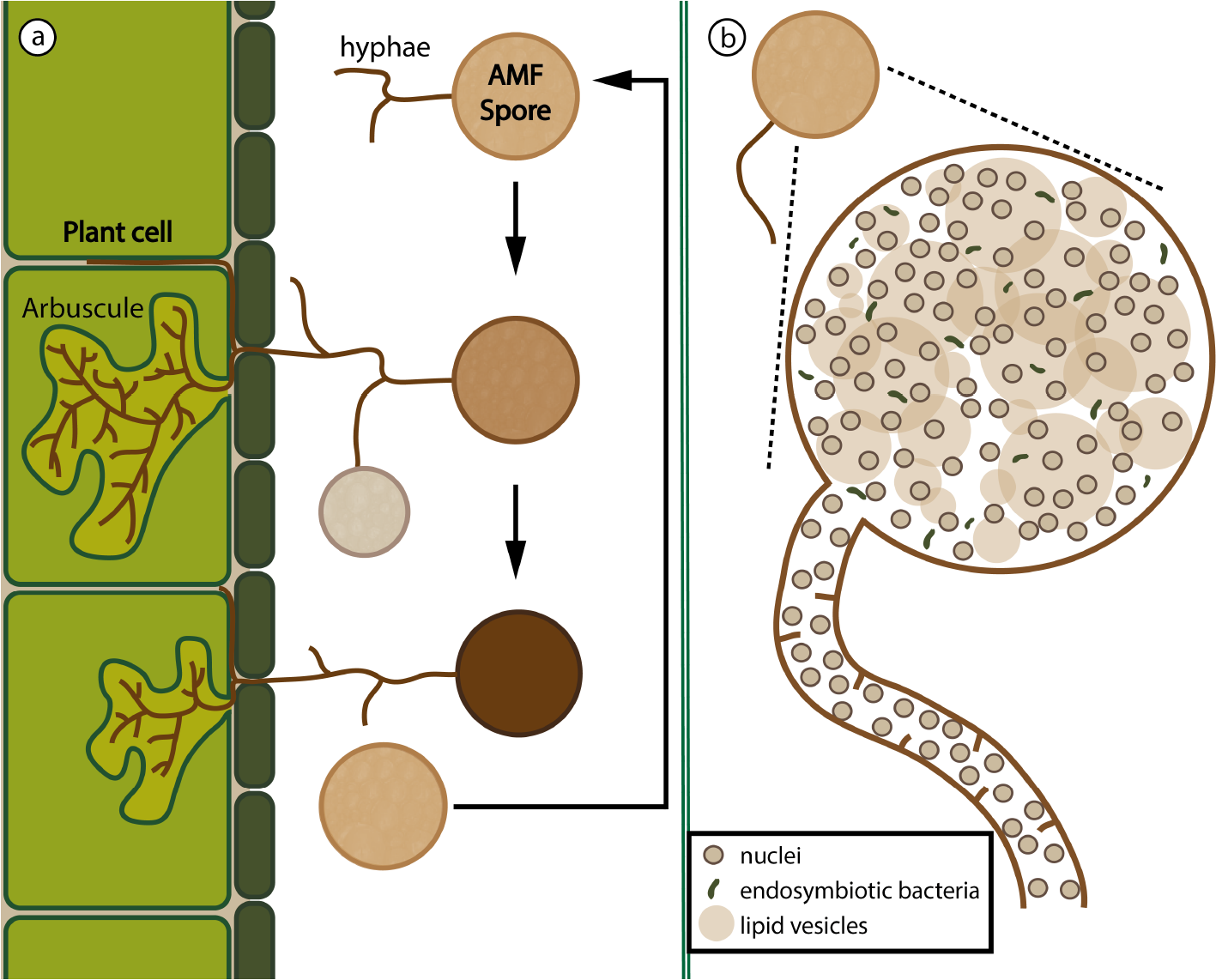
a) Schematic representation of the life-cycle in AM fungi. A spore detects a plant root in the vicinity and grows hyphae towards it. The hyphae penetrate the plant cell wall and form the characteristically branching haustoria with the shape of arbuscules, based on which the group of fungi is named. The arbuscules are used to exchange nutrients with the plant. New spores are produced in other hyphal terminations, bud off upon maturity and remain in dormant state until the cycle starts again, while the first spore dies and the fungi retracts from the plant cell. b) Schematic representation of a spore, containing nuclei, lipid vesicles and endosymbiotic bacteria. The hyphae have very reduced compartmentalization with incomplete septa and nuclei appear to move freely.

A method was developed in which genomic fungal DNA can be obtained, free of plant and microbial DNA, directly from individual nuclei of multinucleate spores. In brief, spores from a trap culture fungal strain of *Claroideoglomus clarodeium/C. luteum* (SA101) were obtained from the INVAM pot culture collection. An initial trial to sort AM nuclei was carried out using pools of spores in order to assess the optimal settings. Cleaned spores were crushed vigorously, and the solution was stained and analyzed by Fluorescence-Activated Cell Sorting (FACS), recording level of fluorescence as a measure of DNA content, and light scattering as proxy for size and particle granularity (Figure 2 a-h). A distinct cloud of particles was observed above the background in the scatter plot (Figure 2h, inside the blue box) which, after microscopy and PCR verification tests with fungal and bacterial specific primers, was confirmed to consist of intact biological structures containing mostly fungal DNA (Figure S1-S3, Table S1). Hence, we concluded that these particles were fungal nuclei and restricted future sorting to this window. Thereafter, individual nuclei from a single spore of the same strain were sorted into wells of a 96-well plate (Figure S4, Table S2) and whole genome amplified (WGA) using multiple displacement amplification (MDA; Figure 2 i-j). The amplified DNA was scored for pure fungal origin by parallel amplification of rDNA barcode regions for both fungi and bacteria (Figure 2k, Figure S5). Twenty-four amplified nuclei samples, confirmed to contain only fungi (Figure S4, Table S3, S4), were sequenced with Illumina HiSeq X (Figure 2l). Further, the MinION Nanopore-based sequencing device (Oxford Nanopore Technologies, ONT, UK) was used to obtain long read sequences for amplified DNA from multiple (5-100) nuclei separated from a pool of 30 spores of the same strain (Figure 2 i-k, m).

**Figure 2.**
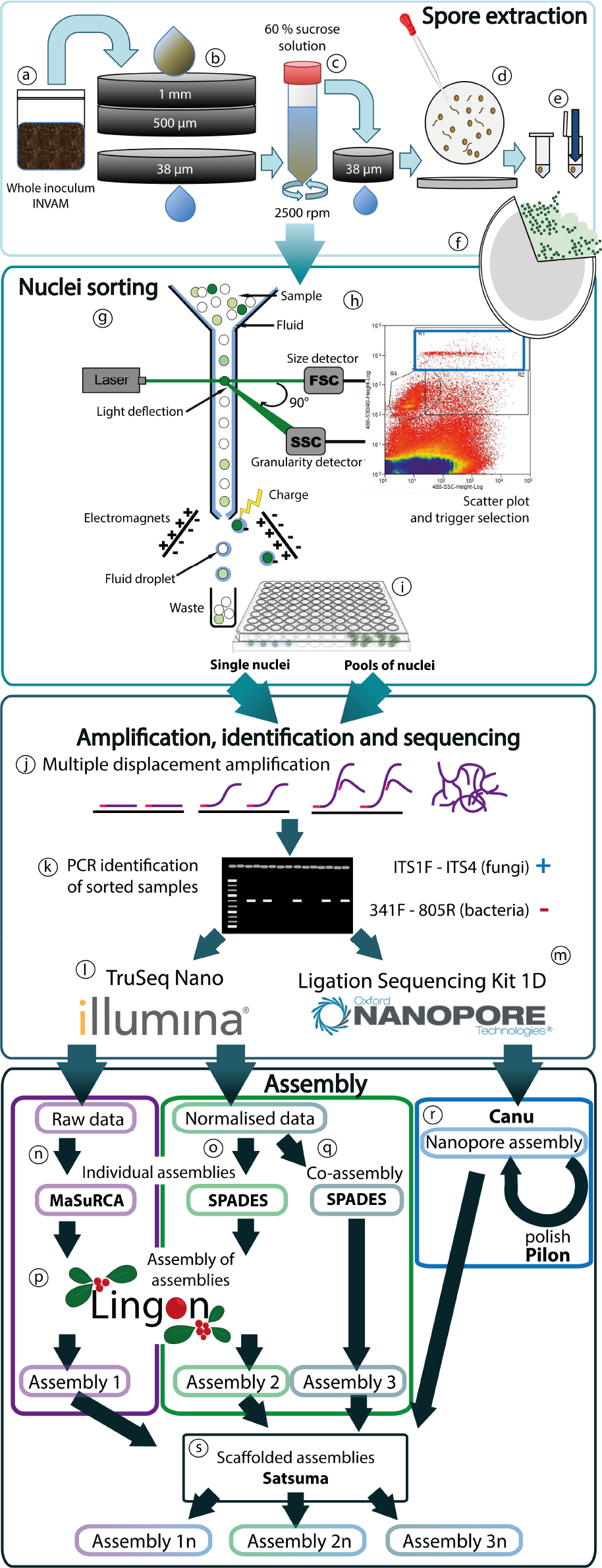
From a soil sample to AM fungal genome assemblies. a) Whole inoculum from the culture collection INVAM is blended with water and (b) poured into a set of sieves, the material stuck in the 38 μm sieve is placed into a (c) tube that contains a solution of 60% sucrose, then centrifuged for 1 min. The supernatant is again run through a 38 μm sieve and washed with water. d) The sieve content is placed in a Petri dish for the spores to be manually picked using a glass pipette. e) After cleaning the spores with ddH_2_O, these are placed one-by-one into tubes and crushed with a pestle. f) The DNA from a broken spore is stained with SYBR Green, giving a strong fluorescent signal for the nuclei, and lighter for the background, organelles and microbes. g) The stained spore content is loaded on the FACS, in which the sample moves inside a constant flow of buffer and crosses a laser beam. An excitation laser of 488-nm and 530/40 band pass filter was used for the SYBR Green fluorescence detection. In addition scattered light, forward scatter (FSC) and side scatter (SSC) were used as proxy for size and granularity to identify the nuclei. h) The signals can be interpreted in a scatter-plot, and particles of a selected cloud (e.g., R1, blue-box) can be sorted individually or pooled (i) into individual wells of a 96-well plate by directing them with a charge. j) The content of each well is whole genome amplified using MDA. k) The amplified products are tested for fungi and bacteria by PCR screening with specific rDNA primers for fungi and bacteria. The products confirmed to be from fungal nuclei are sequenced with l) Illumina HiSeqX, for single nuclei; and m) Oxford Nanopore, for pools of nuclei. To produce assembly 1, Illumina reads are assembled separately for individual nuclei using MaSuRCA^34^ (n). To produce assembly 2, reads are normalized for individual nuclei and assembled with SPADES^35^ (o). For assembly 3 reads from all nuclei are combined before normalization and then assembled with SPADES^35^ (q). Individual nuclei assemblies from method 1 and 2 are assembled together using Lingon^36^ (p). Nanopore data is assembled with Canu^37^ (r), polished with Pilon^38^ using the Illumina raw-reads and used to scaffold the three generated assemblies using Chromosemble, of Satsuma^39^ (s).

Three customized assembly workflows were developed in order to evaluate assembly quality in the light of coverage bias introduced by WGA, which is the biggest challenge when assembling sequence data from amplified single nuclei. The MDA method, however, has an advantage over PCR-based methods in that it produces longer fragments of DNA with a lower error rate, and that the coverage bias is random^40,41^.

For the first two assembly workflows individual nuclei assemblies were generated and subsequently combined to generate a consensus assembly using the workflow manager Lingon^36^ (Figure 2p), which consists of a motif-distance based long sequence overlap finder that merge sequences based on mutual maximal overlaps. In the first assembly workflow, raw Illumina reads were assembled using MaSuRCA^34^ (Figure 2n) resulting in 24 assemblies, ranging in size from 14 to 69 Mbp (Tables S5). In order to overcome MDA generated differences in coverage across the genome the second assembly workflow normalized raw reads to maximum 100X before assembly using SPADES^35^ (Figure 2o), generating 24 assemblies ranging in size from 11 to 50 Mbp (Table S5). A third assembly was created using SPADES^35^ after combining raw reads from 24 nuclei followed by normalization to 100X (Figure 2q). One full assembly with 24 nuclei was generated from each workflow and subsequently scaffolded with a Nanopore assembly built with Canu^37^ (Figure 2r-s). To test for the effect of increasing number of assembled nuclei in the three methods, random combinations with different number of nuclei were assembled with the three assembly workflows. Multiple replicate assemblies were performed for different random combinations of two to twelve nuclei and one random combination for 13-23 nuclei. BUSCO^42^, assembly size and N50 was used to compare these to full and single nuclei assemblies.

## Results

The different assembly workflows resulted in assemblies that vary in sizes, fragmentation and completeness (Table 1). Based on BUSCO analyses, workflow 3 generates the most complete assembly, with 89% for assembly 3n, compared to 2n at 80%, and 1n at 78% (Table 1). Of the core single copy genes identified by BUSCO, few were fragmented or duplicated in assembly 3n indicating that the set of 14,600 predicted genes is likely to be complete and a close representation of the genetic diversity in this strain (Table 1). This number is lower than the number of genes found in other sequenced AM fungi such as *R. irregularis*^28^ and *R. clarus*^31^, also lower than those predicted in assemblies 1n and 2n (Table 1). Interestingly, assembly 3n is considerably smaller (70.8 Mb) than the other full assemblies (92.4 Mb and 130.4 Mb for assembly 1n and 2n respectively), and markedly smaller than the average estimated genome size of 119 Mb based on SGA-PreQC^43^. The smaller size of 3n can be attributed to repeat sequences (20.6 Mb) that are captured, to a lesser extent, compared to the other assembly workflows (41.3 - 58.6 Mb). Specifically, normalization is expected to disproportionally reduce high coverage genomic sequences such as repeat elements, and collapsing those regions when assembling. Note that this effect of normalization is eluded in assembly workflow 2, in which nuclei are normalized and assembled individually; repetitive regions will collapse but in different parts of the genome, ending up represented in the final assembly when combined. In contrast, workflow 1 is based on non-normalized reads. Due to uneven coverage, this workflow assembles less of the genome (an average of 55% of the raw reads align to the individual nuclei assemblies, as opposed to 96% of the reads mapping to the normalized individual nuclei assemblies (Table S5)) but generate contigs well supported by high coverage. Combining these incomplete assemblies from single nuclei using Lingon, generates an accurate assembly 1 comparable to assembly 3 with a better representation of repeats (Table 1).

Combinations of increasing number (1-24) of random nuclei were produced for all the assembly workflows in order to evaluate the number of nuclei needed to produce a good final assembly. As shown in figure 3, single nuclei assemblies are most complete when using normalized workflows (2 and 3), with an average of 40% BUSCO estimated completeness. Interestingly, there is an increasing number of gene duplications among the complete genes as more single nuclei assemblies are combined for method 2 compared to method 1 (Figure 3a-b). Higher amount of gene duplications was confirmed by locating known single copy genes in all assemblies (Table S6). The duplications in workflow 2 are likely generated because read normalization allows for assembly of regions with low coverage that are prone to errors and prevents contigs from being properly assembled by the workflow manager Lingon.

## Discussion

Methodological challenges in assembling genomes from amplified single nuclei or cells can be elevated by careful analysis of generated assemblies. Combining and normalizing reads (workflow 3) from only 6 individually sequenced nuclei can already generate a high coverage genome assembly. From this assembly, good quality data of single copy genes are obtained, ideally suited for phylogenomic studies. Assembly workflow 1 on the other hand is better suited to characterize repeat elements in the genome since these are better represented in assemblies of non-normalized data. With this method a high-quality genome can be assembled using seven individually amplified and sequenced nuclei (Figure 3). Comparative genetic analysis of single nuclei is best done using assemblies from workflow 2. However, single nuclei assemblies based on normalized reads should not be assembled into consensus assemblies since variable quality of contigs make them prone to duplication.

To conclude, sequence data from single cell sequencing presents itself as challenging, but as shown here, with the right combination of methods adapted to the data, *de novo* genome references can be generated, opening the door for an expansion in genomic and phylogenomic research in organisms like AM fungi, that have, for too long, evaded large scale genome sequencing efforts due too methodological limitations stemming from their complicated biology. Useful genomic information can be generated from a handful of single nuclei greatly improving our ability to study multicellular eukaryotes with complex life stages. The assembly method of choice will ultimately depend on the research questions asked and the kind of data needed or available.

**Table 1.**
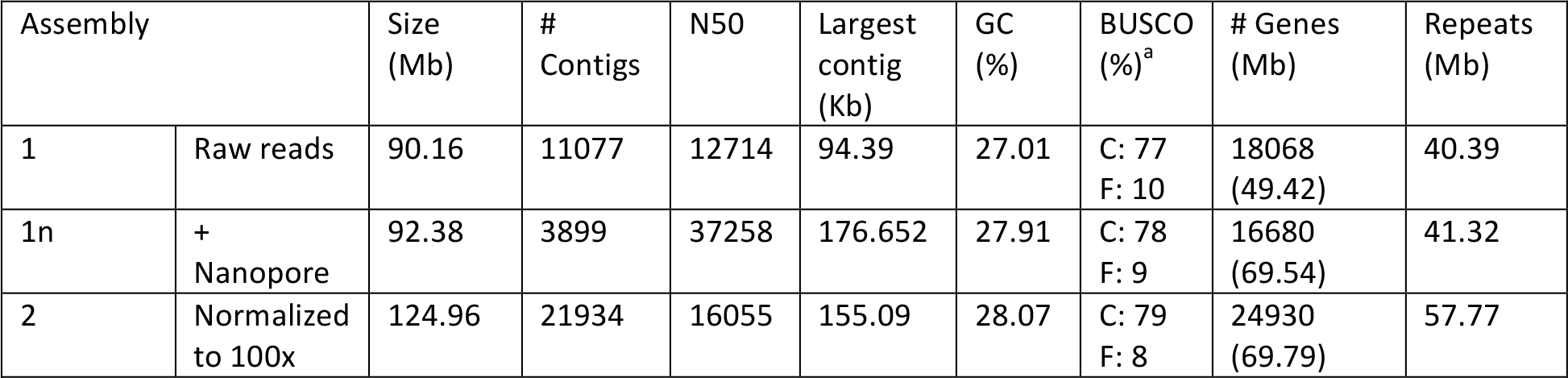
Comparative assessment of the 3 assembly workflows.

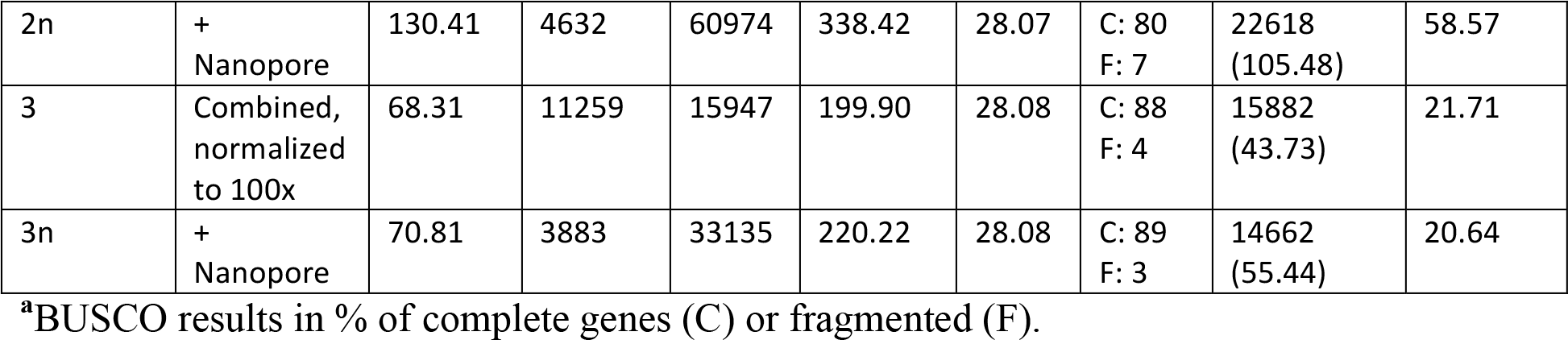

**Figure 3.**
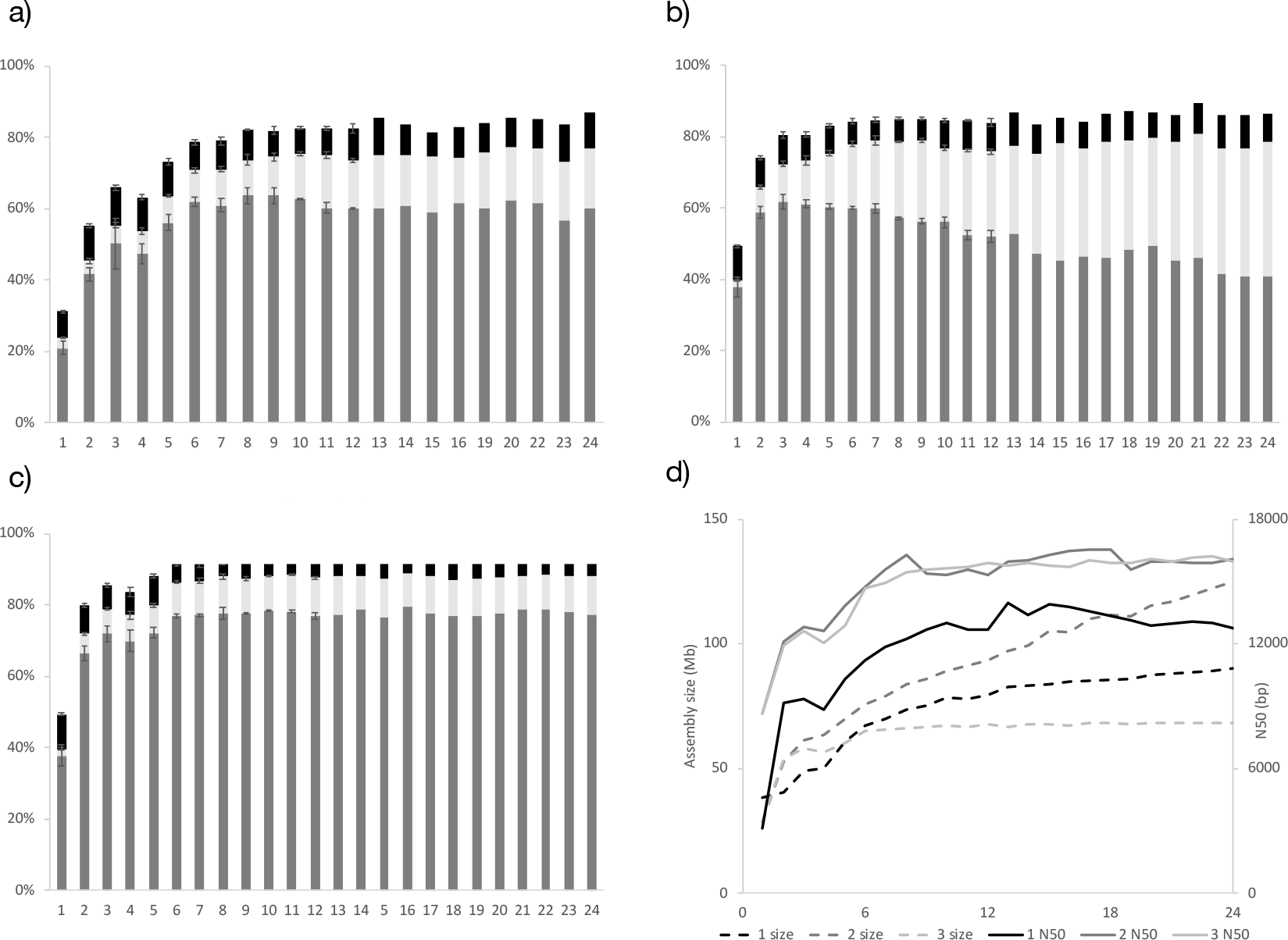
Summary statistics for different number of assembled nuclei (1-24) using three different assembly workflows. BUSCO estimates of completeness for a) workflow 1: raw reads of individual nuclei assembled using Masurca, consensus assembly using Lingon b) workflow 2: normalised reads of individual nuclei assembled using SPADES, consensus assembly using Lingon and c) workflow 3: reads from individual nuclei are pooled and normalised before assembling with SPADES. Percentage of single copy core genes detected as single copy (S: grey), duplicated (D: light grey) or fragmented (F: black). Average of 3-6 replicate assemblies up to 12 nuclei with error bars indicating SEM. In d) assembly size (dashed lines) and N50 (solid lines) for the there methods 1 (black), 2 (grey) and 3 (light grey).

## Methods

### Fungal strain and spore extraction

*C. claroideium/C. luteum* (SA101) was obtained as whole inoculum from the International culture collection of (vesicular) arbuscular mycorrhizal fungi (INVAM) at West Virginia University, Morgantown, WV, USA. Soil (10-30 ml) was blended with 3 to 4 pulses using a blender half-filled with water (500 ml). The mix was filtered through a set of sieves (1 mm/500 μm/38 μm × 200 mm diameter, (VWR, Sweden). The content of the last sieve was transferred into a falcon tube containing 20 ml of 60% sucrose solution, and centrifuged for 1 minute at 2500-3000 rpm. The supernatant was poured into a small sieve (50 mm diameter) of 38 μm and the sucrose was washed with water. The contents were poured onto a petri dish for better visualization under the stereomicroscope. Spores were transferred individually or in groups to an Eppendorf tube using modified glass pipettes with reduced tip diameter and subsequently cleaned by adding and removing ddH2O five times. The step-by-step protocol can be found in the OSF Repository for the project^44^.

### Nuclei extraction and sorting

After spore extraction from soil, individual spores were placed in 30 μl ddH2O in 1.5 ml Eppendorf tubes. One tube with 15 spores was used to establish the sorting window. An amount of 50 μl 1x PBS was added to each tube before crushing the spores using a sterile pestle. DNA was stained by adding 1 μl of 200x SYBR Green I Nucleic Acid stain (Invitrogen™, Thermo Fisher Scientific, MA, USA) and the sample was incubated for 20-50 min in the dark. More 1x PBS was added to increase the volume to 100-200 μl before putting the sample on the FACS. The sorting was performed with a MoFlo™ Astrios EQ sorter (Beckman Coulter, USA) using a 488 nm laser for excitation, 70 μm nozzle, sheath pressure of 60 psi, and 0.1 μm filtered 1x PBS as sheath fluid. The sorter was triggered on forward scatter (FSC) at a threshold of 0.03% and sort regions were set on SYBR Green I fluorescence (488-530/40) over side scatter (SSC). The samples were sorted in enrich mode with a drop envelope of 1 at 700 to 1200 events per second. Thus, if a particle fitting within the sorting window passes by the laser together with another particle, these would be discarded. Particles from region R1, assumed to be nuclei (Figure S4), were sorted individually into 96 well plates containing 1 μl 1x PBS/well, groups of 5 particles were collected for positive control, and empty wells were kept as negative control (Table S2).

### Whole Genome Amplification

Sorted nuclei were lysed and neutralized followed by whole genome amplification using Phi29 and MDA as described by Rinke et al., 2014^45^. In short, the cells were incubated in an alkaline solution (buffer DLB and DTT, Qiagen, Germany) for 5 min at room temperature, followed by 10 min on ice. Lysis reactions were neutralized by adding 1 μL neutralization buffer (stop solution, Qiagen, Germany). Both, the alkaline lysis solution as well as the neutralization buffer were UV treated with 2 Joule in a Biolinker. MDA was performed using the RepliPHI™Phi29 Reagent set (RH031110, Epicenter, WI USA) at 30°C for 16 h in 15 μl reaction volumes with a final concentration of 1x reaction buffer, 0.4 mM dNTPs, 10 mM DTT, 5% DMSO, 50 μM hexamers with 3’-phosphorothioate modifications (IDT Integrated DNA Technologies, Iowa USA), 40 U Phi 29 enzyme; 0.5 μM SYTO13® (InvitrogenTM, Thermo Fisher Scientific, MA, USA) and water. All reagents except SYTO13 were UV decontaminated with 3 Joule in a UV crosslinker as described in Rinke et al., 2014^45^ 12 μl of MDA mix were then added to each well.

The whole genome amplification was monitored in real time by detection of SYTO13 fluorescence every 15 minutes for 16 h using a Chromo4 real-time PCR instrument (Bio-Rad, USA) or a FLUOstar®Omega plate reader (BMG Labtech, Germany). The single amplified genome DNA was stored at −20°C for short-term, and transferred to −80°C for long-term storage.

### Selecting single amplified nuclei for sequencing

MDA products were diluted to approximately 5 ng/μl (40x) and screened for the presence of fungal and bacterial ribosomal genes using PCR. Reaction mixtures were made as described above, using the fungal-specific primers ITS9^46^ and ITS4. The PCR protocol had an initial denaturing step of 10 min at 95°C, followed by 35 cycles of 30 s at 95°C, 30 s at 58°C, and 50 s at 72°C for the fungi PCR. For the bacteria-specific 341F/805R^47^ primer pairs a different reaction mixture was used containing 10x Standard Taq Reaction buffer (Qiagen), 2 mM MgCl2, 0.2 mM deoxynucleoside triphosphates (dNTPs), a 0.2 μM concentration of each primer, and 1 U Taq DNA polymerase (Qiagen). A positive control of DNA extracted from commercially available *Agaricus bisporus* provided by Dr. Ylva Strid, UU, was included, and ddH2O as negative control. The bacterial PCR protocol consisted on an initial step of 5 min at 95°C, followed by 30 cycles of 30 s at 95°C, 30 s at 58°C, and 50 s at 72°C for the bacteria PCR before a final elongation step of 7 min at 72°C. Bacteria PCR included a positive control of DNA extracted from Legionella provided by Tiscar Graells, Universitat Autónoma de Barcelona (UAB), Spain, and ddH2O as negative control. The reaction was performed with a 2720 Thermocycler of Applied Biosystems (USA). The presence of amplification products was verified by gel electrophoresis by separation on a 2% agarose gel run for 35 min at 110V (fungi) and 70V (bacteria) including a Thermo Scientific GeneRuler 100 bp DNA Ladder. (Figure S5), and the samples were identified as fungi positive, bacteria positive, fungi + bacteria positive or failed/empty (Table S3). From the samples that scored positive for presence of fungi, 24 undiluted samples were selected for sequencing and the DNA amount was measured using Qubit (Brand, country) after addition of 30 μl ddH2O (Table S4).

### Sequencing of single amplified nuclei

From the 24 selected samples, around 800 ng of DNA was transferred to sequencing plates. Library preparation and sequencing was performed by the SNP&SEQ Technology Platform in Uppsala at the National Genomics Infrastructure (NGI) Sweden and Science for Life Laboratory. For each sample, an individual library was prepared using the TruSeq Nano DNA Library Prep Kit. The sequencing was performed by doing a cluster generation and 150 cycles paired-end sequencing of the 24 libraries in 1 lane using the HiSeq X system and v2.5 sequencing chemistry (Illumina Inc., USA). Read data were delivered to us as fastq.

### Spore sorting for Nanopore sequencing

Spores were picked in groups of 30 with the help of a P10 and P100 pipette, then washed 5x in nuclease-free water and transferred to Eppendorf tubes in 30 uL nuclease-free water. For the FACS sorting, spores were crushed, then 30 μl 1x PBS was added to the tube along with 1 μl of 200x SYBR Green for staining the DNA (20-50 mins). Sample volume was increased to 200 μl with 1x PBS before loading on the FACS. Pools of 5 and 100 nuclei were sorted into either individual 1.5 ml Eppendorf tubes or into multi-well plates. The above-described WGA protocol was run, and the presence of fungal DNA in the samples was verified by PCR on diluted samples of amplified pooled nuclei before selecting fungi positive samples for library preparation. PCR reaction mixtures contained 10x Standard Taq Reaction buffer (Qiagen), 2 mM MgCl2, 0.2 mM deoxynucleoside triphosphates (dNTPs), a 0.2 μM concentration of each primer, and 1 U Taq DNA polymerase (Qiagen). The fungal-specific ITS1F/ITS4 and bacteria-specific 341F/805R primer pairs were used for each sample in two independent PCR reactions. The PCR protocol included an initial denaturing step of 5 min at 95°C, followed by either 35 cycles of 30 s at 95°C, 30 s at 55°C, and 50 s at 72°C for the fungi PCR or by 30 cycles of 30 s at 95°C, 30 s at 58°C, and 50 s at 72°C for the bacteria PCR before a final elongation step of 7 min at 72°C. The reaction was performed with a 2720 Thermocycler of Applied Biosystems (USA). Amplification products were visualized and documented by gel electrophoresis as described above.

Libraries were prepared by following the “Premium Whole Genome Amplification” protocol (version WAL_9030_v108_revJ_26Jan2017, Oxford Nanopore Technologies [ONT], Oxford, United Kingdom) in combination with the Ligation Sequencing Kit 1D (SQK-LSK108, ONT) with the following modifications: (a) an alternative WGA method was used (Qiagen Single Cell Kit instead of the Midi Kit); (b) samples were diluted to a 50 μl volume following WGA and quantified with a Qubit fluorometer (brand, country). Amounts of 1 - 2.5 μg DNA were then used for preparing individual libraries, starting with the first bead cleaning step explained in the whole genome amplification section. At the end of this step, samples were eluted in 19 μl nuclease-free water instead of 100 μl. 1 μl of the eluted sample was used for DNA quantification (Qubit fluorometer) while another 1 μl was used to measure DNA quality with Nanodrop (ND 2000); (c) no size selection and intentional shearing was performed to achieve read length as long as possible; (d) 17 μl amplified DNA was added to the T7 endonuclease treatment; (e) an extended end-prep reaction was performed by incubating the samples for 30-30 mins at both 20°C and 65°C; (f) adapter ligation was allowed for 25-30 mins instead of 10; (g) elution buffer in the final step was incubated for 15 minutes instead of 10; (h) the loaded library contained no additional water but 14.5 μl DNA library instead of 12 μl. Additionally, flicking was used to mix reactions instead of pipetting to prevent DNA fragmentation. Further, eluates were removed and retained in a stepwise fashion (i.e. in multiple aliquots) after every cleaning step to assure that no beads were brought forward with the DNA into the next library preparation step. In general, by extending clean-up-, ligation- and elution steps the quality of the library and thus pore occupancy during sequencing could be improved.

A total of 3 libraries on 3 separate ONT MinION R9.4 flow cells (FLO-MIN106) were sequenced using live base-calling and the standard 48 h sequencing protocol (NC_48Hr_sequencing_FLO-MIN106_LSK-108_plus_Basecaller). One library was run on a fresh flow cell with ~1400 single pores available for sequencing in the beginning of the run. This 48 h run provided 1,686,715 reads. As for the other two libraries, previously used and washed flow cells were re-used with only a fraction of sequencing pores being functional (402 vs. 256 pores), thus the acquired data were much lower (100,000 and 106,000 reads respectively).

### Computational analysis, assembly and annotation

The quality of the Illumina reads was assessed with FastQC^48^. Genome size estimation was done for each paired raw-reads from individual nuclei with SGA-PreQC^49^. Contamination was assessed with Kraken^50^ in some of the raw-reads. CG content was computed using the NBIS-UtilityCode^51^ toolbox.

Assembly workflow 1: Individual assemblies for each of the 24 nuclei was done using MaSuRCA^34^ using default options. The resulting assemblies were iteratively merged using Lingon^36^, which computed overlaps based on the spacing of sequence motifs (CATG, CTAG, GTAC, GATC, TATA, ATAT, and GC), and merged contigs based on pairwise maximal extensions. Each motif was iterated over ten times. Three versions of the assembly were generated when contigs smaller than <500, <1000 and <2000 were removed from the individual assemblies prior to Lingon.

Assembly workflow 2: Each set of reads was normalized using bbnorm of BBMap^52^ v. 38.08 with a target average depth of 100x. Normalized data were assembled individually into 24 assemblies using SPADES^35^, and a consensus assembly was generated with Lingon^36^, with the same sequence motifs as for assembly 1.

Assembly workflow 3: The 24 datasets were combined and normalized with bbnorm of BBMap^52^ v. 38.08 with a target average depth of 100x, and posteriorly assembled using SPADES^35^.

Nanopore assembly: Nanopore reads were assembled using Canu^37^ v.1.7-86da76b, this specific beta version made it possible to assemble a difficult dataset like ours, with highly uneven coverage across the genome. An assembly was created using default settings together with the known information (genomeSize=117m -Nanopore-raw). The resulting individual assembly was polished with three rounds of Pilon^38^ v.1.22 using the raw Illumina reads from the 24 nuclei mapped with Bowtie2^53^. The contigs of the final assemblies from single nuclei were scaffolded with the Nanopore assembly using Chromosemble from the Satsuma package^39^.

### Comparative assembly analysis

A quantitative assessment of the assemblies was done with Quast^54^ v.4.5.4 and a contamination check with Kraken^50^ v1.0. In addition, a BUSCO^42^ analysis was done to assess completeness of the genome. The BUSCO lineage set used was fungi_odb9 and the species set was rhizopus_oryzae. (Figure 3, Figure S Raw-reads were mapped to the individual assemblies of method 1 and 2 (Table S5) with Bowtie2^53^ v. 2.3.3.1 using the default settings.

Two genes, known to be single copy genes in fungal genomes, as elongation factor 1-alpha (EF1-alpha) and the largest subunit of RNA polymerase II (RPB1), were searched for in the genome assemblies to test for possible duplications generated by the assembly methods. Sequences belonging to *C. claroideum* were used to find the sequences with BLASTn^55^ (Table S6). Genebank sequences: EF1-alpha GQ205008.1, RPB1 HG316018.1.

### Genome annotation

Repeats and transposable elements (TEs) were *de novo* predicted in every assembly using RepeatModeler^56^ v1.0.8. The repeat library from RepeatModeler was used to mask the genome assembly using RepeatMasker^57^ v4.0.7. The classification reports can be found in the OSF Repository^44^.

Protein coding genes were *de novo* predicted from the repeat-masked scaffolded genome assembly with GeneMark-ES^58^ v4.33. GeneMark-ES uses unsupervised self-training and an algorithm that is optimized for fungal gene organization. To guide the gene predictions, we aligned UniProt/Swiss-Prot^59^ protein sequences (downloaded 8 May 2018) to the repeat-masked genome assembly using MAKER^60^ v3.01.1-beta and provided the genomic locations of the protein alignments to GeneMark-ES. The previously published transcriptomic data from *C. claroideum*^61^ was not used to due to the low mapping success of the reads to the assembly (25%), which could be related to the low BUSCO statistics shown in the study^61^, and that could have negatively affected the annotation quality.

Protein and gene names were assigned to the gene predictions using a BLASTx^55^ v2.6.0 search of predicted mRNAs against the UniProt/Swiss-Prot^59^ database with default e-value parameters (1×10-5). The ANNotation Information Extractor, Annie^62^, was used to extract BLAST matches and to reconcile them with the gene predictions.

Sequences, assemblies and annotation can be found in the BioProject: PRJNA528883

## Acknowledgments

We thank, J. Bever and S. Bertilsson for scientific discussions, Y. Strid and M. Zakieh for assistance in the lab, J. Morton and W. Wheeler at INVAM culture collection, and funding from ERC (678792). Sequencing was performed by the SNP&SEQ Technology Platform at NGI Sweden and SciLife Laboratory, Uppsala, supported by the VR and the KAW. Computations were performed on resources provided by SNIC through UPPMAX.

## Author contributions

AR initiated the project and developed the method together with MMN and HJ. MSG helped develop the bioinformatic analysis. CB was in charge of the single nuclei facility and did the MDA, BE was in charge of the Nanopore sequencing. MG designed Lingon and helped with analysis together with MK. VK was in charge of the annotation. MMN was responsible for the project and wrote the manuscript with AR and HJ, with with input from all the authors.

