## Supplementary Information for "From single nuclei to whole genome assemblies"

### Developing the protocol of single nuclei extraction

Broken spores were observed in a microscope equipped with a fluorescent filter to assess amount, release efficiency and size of the nuclei. Spores were placed on a slide and the excess water was removed with a pipette. A small drop of Gold Slow Fade + DAPI was added and the glass cover was placed pressing gently with the back of a pencil, to crush open the spore without destroying it. Crushed spores were visualized under fluorescent microscope. (Figure S1, Original pictures in OSF Repository[44]).

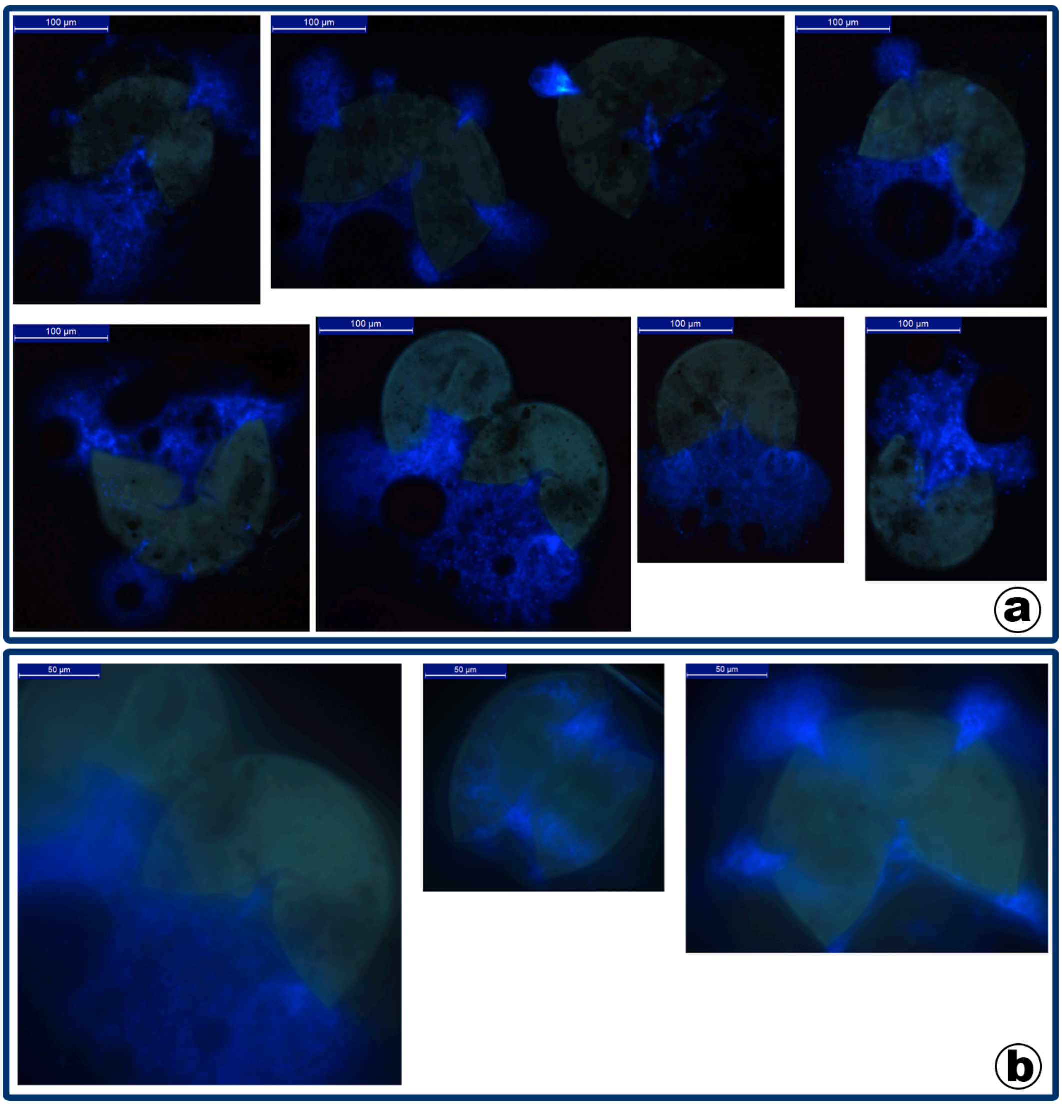

Figure S1. Spores of *C. clarodeium/C. luteum* (SA101) at 20X (a) and 40X (b) seen in the microscope after staining with DAPI in order to visualize the nuclei and assess their size.

For the sorting of the nuclei, two tubes with 15 and 25 spores of the isolate respectively, previously extracted and cleaned were crushed in 200 µl of ddH2O, and 200 µl of 1X PBS was added together with 1.25 µl of 200X SYBR Green. After 37 min staining, 100 µl of 1X PBS was added to increase the volume before running the sample in the FACS. FSC on the 488 nm laser was used as trigger, and a 530±20 nm (530/40) band pass filter for fluorescence signal detection. The best resolution of distinctive populations of particles in the scatter plot was obtained when plotting level of fluorescence against the SSC representing granularity (Figure S2).

Three regions were defined in the scatter plot: R1 (a distinctive line with strong fluorescence), R4 (a distinctive community lower fluorescence and lower SSC values than R1), and R2 minus R4 (an undifferentiated area between R1 and the background fluorescence) (Figure S2). Particles appearing in those areas were sorted simultaneously into three different tubes (Table S1). Additionally, two tubes were filled with particles from all the areas in each sorting round, to be used as a positive control.

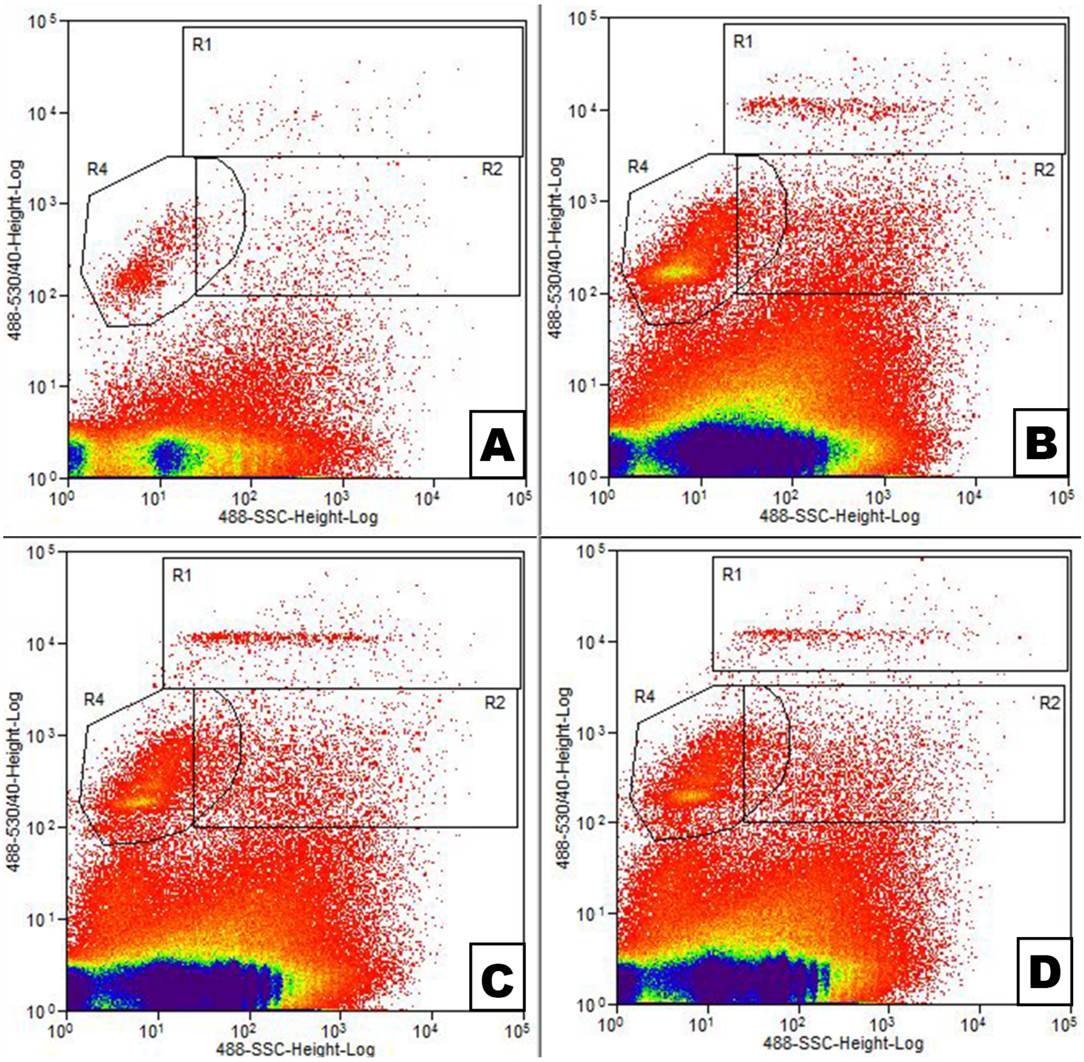

Figure S2. Scatter plots from the FACS sorting of particles from 15 (a, b) or 25 (c, d) crushed spores using Forward Scatter (FSC) as trigger. On the x axis, SSC, on the y axis, the SYBR Green fluorescence. The low-fluorescence part of the scatter plots represents background fluorescence in the spore content, that is either not stained or very lightly stained. a) Scatter plot of from the first quick run identifying the areas to sort from. b) Scatter plot of actual sorting into pools (R1, R2, R4) of particles from the 15 spores (Table S1). c) Scatter plot of sorting into pools (R1, R2, R4) of particles from the 25 spores (TableS1). d) Scatter plot when sorting only the area R1 from 25 crushed spores (Table S1).

DNA was extracted from each of the tubes using the Genomic DNA from Plant kit (Macherey-Nagel, Germany) in order to obtain clean DNA that could be tested for fungal presence. The protocol was adapted to the characteristics of the samples having low amounts of material and no cell biological material to homogenize, these encompass the following: step 1, 125 µl of buffer MC1 and 2.5 µl of RNAse A; step 2 was eliminated, but a short centrifuge of 30 s at 5800 xg was added to bring down the evaporates; step 3, 7.5 µl magnetic beads and 100 µl MC2; step 4, 140 µl MC3; step 5, 140 µl MC4; step 6, 125 µl of 80% ethanol; step 7, 125 µl MC5; step 8, 30 µl MC6. Nanodrop was used to determine the amount of DNA in the samples after the DNA extraction (Table S1).

Table S1. Amount of particles collected for each region and assessment of the DNA amount using nanodrop after extracting DNA from the samples.

| Sample | | Number of particles | Nanodrop (ng/µl DNA) |
|---|---|---|---|
| 15 spores | R4 | 10795 | 21.5 |
| | R1 | 778 | 22.7 |
| | R2 (- R4) | 4846 | 28.1 |
| | All regions | - | 20.8 |
| 25 spores | R4 | 16000 | 26.0 |
| | R1 | 1300 | 28.2 |
| | R2 (- R4) | 8000 | 22.5 |
| | R1 | 622 | 26.4 |
| | All regions | - | 25.6 |

Finally, to resolve in which of the regions of the scatter plot the nuclei might be falling into, PCR was done using the fungal specific ITS1F/ITS4 primers[63] for amplification of the ribosomal RNA region. Reaction mixture contained 5x Buffer HF (Qiagen, Sweden), 2 mM MgCl2, 0.2 mM deoxynucleoside triphosphates (dNTPs), 0.4 mM DMSO, a 0.25 μM concentration of each primer, and 1 U Phusion DNA polymerase (Qiagen, Sweden). The PCR protocol included an initial denaturing step of 30 s at 98°C, followed by 35 cycles of 30 s at 98°C, 30 s at 57.5°C, and 30 s at 72°C, before a final elongation step of 10 min at 72°C. The reaction was performed with a 2720 Thermocycler of Applied Biosystems (Fishers Scientific, Sweden). Amplification products were separated by gel electrophoresis (1.5 % agarose) at 70V for 80 min (details) and visualized using gel doc system (details). From the visualization we could resolve that fungal nuclei were most frequently appearing in the region R1 of the scatter plot, as these samples generally showed stronger amplification of the fungal barcode region (Figure S3).

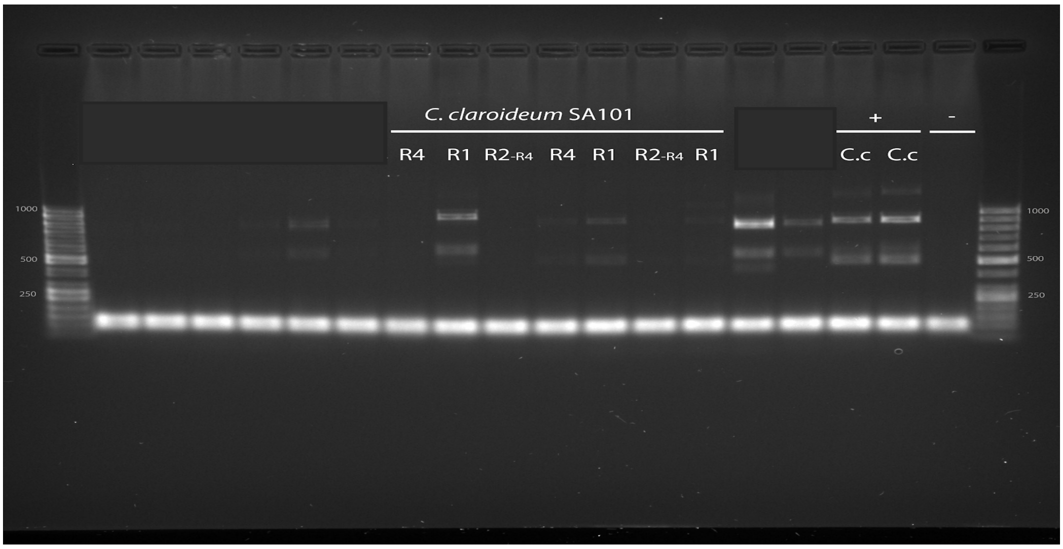

Figure S3. Agarose gel showing the PCR products for particles collected from the scatter plot regions R1, R2-R4, R4 as well as a pool of all regions as positive control sorted from *C. clarodeium/C. luteum* (SA101). Gel electrophoresis was done using a

1.5% agarose gel, run for 80 min at 70 V. Thermo Scientific GeneRuler 50 bp DNA Ladder was used.

**Nuclei extraction and sorting**

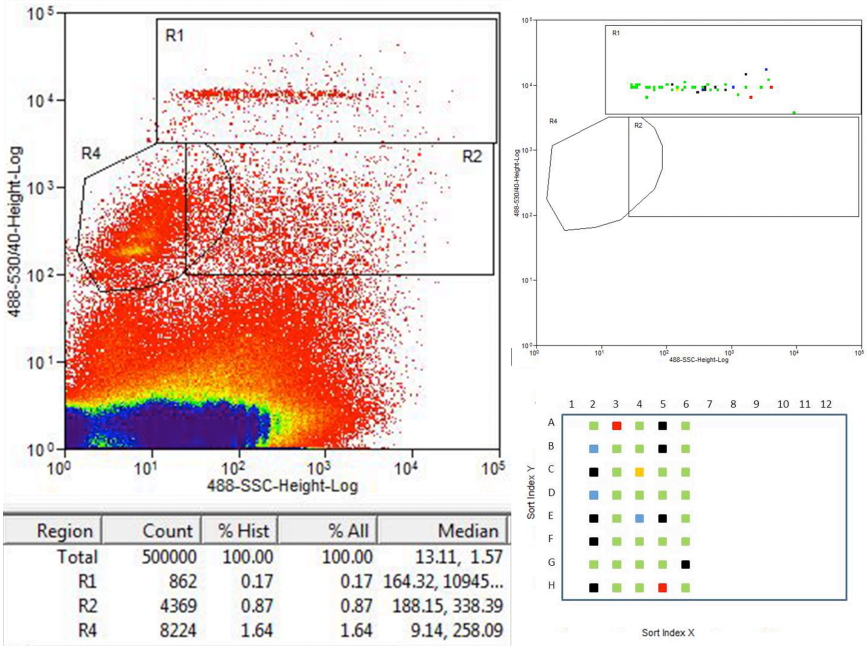

Figure S4. On the left, scatter plot for one single spore crushed from which the R1 region was sorted. On the right, the particles that were sorted and were further tested for fungi/bacteria. Every particle can be traced back from the plate to the exact sorting event. Colors represent the obtained results observed in Figure S5: fungal positives/bacteria negatives (green), fungal positives/bacteria positives (blue), fungal negatives/bacteria positives (red), no amplification (black).

Table S2. 96 well plate layout for sorting. In blue, empty wells as negative controls; in light orange, single particles; in dark orange, 5 particles for positive control. Half plates were filled to reduce risk and cost in downstream handling.

| | 1 | 2 | 3 | 4 | 5 | 6 | 7 | 8 | 9 | 10 | 11 | 12 |
|---|---|---|---|---|---|---|---|---|---|---|---|---|
| A | | 1 | 1 | 1 | 1 | 1 | | | | | | |

| | | | | | | |
|---|---|---|---|---|---|---|
| B | | 1 | 1 | 1 | 1 | 1 |
| C | | 1 | 1 | 1 | 1 | 1 |
| D | | 1 | 1 | 1 | 1 | 5 |
| E | | 1 | 1 | 1 | 1 | 1 |
| F | | 1 | 1 | 1 | 1 | 1 |
| G | | 1 | 1 | 1 | 1 | 1 |
| H | | 1 | 1 | 1 | 1 | 5 |

**Selecting single amplified nuclei for sequencing**

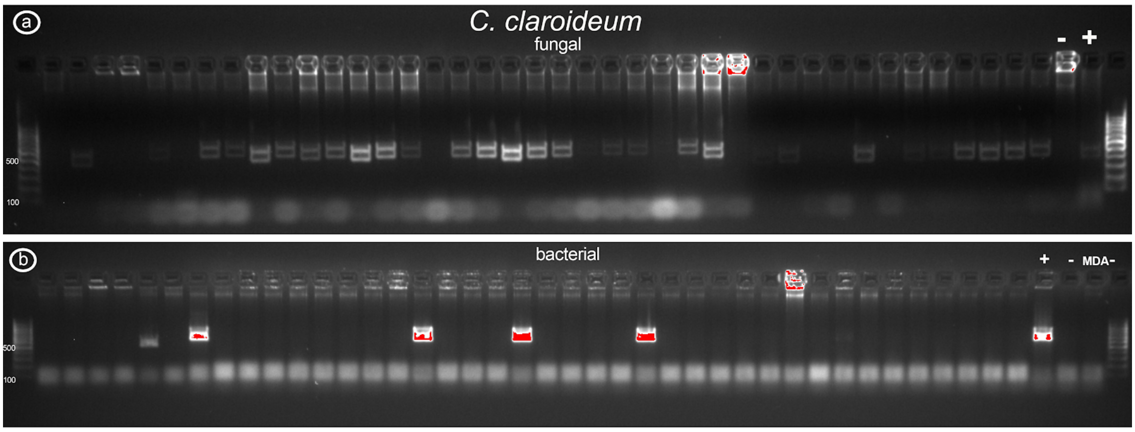

Figure S5. PCR test on the sorted and amplified single nuclei samples for a) Fungi (Positive control *Agaricus bisporus,* negative control ddH2O). b) Bacterial (positive control *Legionella,* negative control ddH2O) . Gel electrophoresis was done using a 2% agarose gel, run for 35 min at 110 V and 70 V respectively. Both ladders are Thermo Scientific GeneRuler 100 bp DNA Ladder.

Table S3. Total number (percentage of total) of sorted and amplified nuclei samples falling into the four categories Fungi +, Bacteria +, Fungi and Bacteria + and Failed/Empty, based on PCR scoring described above. Total number sorted and tested in the last column.

| Species | Fungi + | Bacteria + | Fungi and Bacteria + | Failed/Empty | Total sorted |
|---|---|---|---|---|---|

| | | | | | |
|---|---|---|---|---|---|
| *C. clarodeium/C. luteum* (SA101) | 27 (67.5%) | 2 (5%) | 3 (7.5%) | 8 (20%) | 40 |

Table S4. DNA concentrations measured with Qubit on the 24 samples of whole genome amplified DNA from sorted nuclei of *C. clarodeium/C. luteum* (SA101)selected for sequencing.

| Sample | Well code | DNA concentration (ng/µl) |
|---|---|---|
| 1 | A2 | 24.5 |
| 2 | B2 | 26.0 |
| 3 | D2 | 22.5 |
| 4 | C3 | 23.3 |
| 5 | D3 | 23.0 |
| 6 | F3 | 23.2 |
| 7 | G3 | 23.8 |
| 8 | H3 | 24.9 |
| 9 | A4 | 22.2 |
| 10 | B4 | 23.8 |
| 11 | C4 | 23.7 |
| 12 | D4 | 26.2 |
| 13 | E4 | 22.6 |
| 14 | F4 | 26.5 |
| 15 | G4 | 21.7 |
| 16 | H4 | 23.7 |
| 17 | C5 | 22.6 |
| 18 | D5 | 26.4 |
| 19 | G5 | 23.2 |
| 20 | A6 | 27.2 |
| 21 | B6 | 21.5 |
| 22 | D6 | 23.6 |

| | | |
|---|---|---|
| 23 | E6 | 21.4 |
| 24 | F6 | 23.8 |

**Quality control**

In the OSF repository for the project[44] all quality control results for the raw reads are deposited, as well as the qualitative and quantitative assessments of the assemblies. The standard of Minimum Information about a Single Amplified Genome (MISAG) developed by the Genomic Standards Consortium (GSC)[64] was used as a guideline to assess the quality of the assemblies (Table S5). The highest quality reached in this study is the high-quality draft, considering that the final assembly contains still gaps spanning repetitive regions.

Table S5.

Single nuclei assembly statistics for the assembly workflow 1 (raw reads assembled with MaSuRCA[34]) and 2 (normalized reads assembled with SPADES[35]). Assembly size (Mb), number of contigs, N50, size of largest contig and % of raw reads covered by the individual assembly is presented for each nucleus (1-24).

| | Assembly workflow 1 (raw reads + MaSuRCA) | | | | | Assembly workflow 2 (normalized reads + SPADES) | | | | |
|---|---|---|---|---|---|---|---|---|---|---|
| nucleus | Size (Mb) | # Contigs | N50 | Largest contig (Kb) | % or raw reads mapped | Size (Mb) | # Contigs | N50 | Largest contig (Kb) | % or raw reads mapped |
| 1 | 35.44 | 16406 | 3231 | 36.3 | 53.44 | 27.9 | 7592 | 8568 | 58.9 | 95.52 |
| 2 | 31.29 | 15557 | 2959 | 33.1 | 54.11 | 24.90 | 6609 | 9212 | 199.9 | 96.81 |
| 3 | 25.03 | 16526 | 1672 | 13.2 | 50.77 | 20.14 | 7764 | 5294 | 61.2 | 95.99 |
| 4 | 31.10 | 14480 | 3149 | 25.6 | 52.36 | 22.98 | 6175 | 8841 | 62.4 | 95.66 |
| 5 | 39.00 | 17793 | 3351 | 32.9 | 55.70 | 27.72 | 7429 | 9343 | 78.0 | 95.29 |
| 6 | 36.19 | 16212 | 3510 | 46.6 | 51.94 | 27.38 | 7453 | 9824 | 63.8 | 95.87 |

| | | | | | | | | | | |
|---|---|---|---|---|---|---|---|---|---|---|
| 7 | 56.06 | 26206 | 3199 | 31.9 | 52.88 | 41.98 | 9863 | 10068 | 95.0 | 95.82 |
| 8 | 51.87 | 23590 | 3237 | 38.9 | 51.08 | 39.23 | 9291 | 10447 | 60.6 | 95.83 |
| 9 | 42.24 | 19220 | 3284 | 30.1 | 52.42 | 30.72 | 8355 | 9500 | 71.7 | 95.86 |
| 10 | 51.87 | 23590 | 3237 | 38.9 | 61.59 | 16.74 | 5150 | 7065 | 55.5 | 96.11 |
| 11 | 14.47 | 7425 | 2705 | 22.5 | 48.11 | 11.78 | 3500 | 7214 | 41.2 | 96.31 |
| 12 | 45.21 | 21122 | 3120 | 42.3 | 53.31 | 33.37 | 8415 | 9772 | 74.8 | 95.74 |
| 13 | 56.85 | 24537 | 3785 | 38.2 | 60.21 | 40.47 | 10218 | 11172 | 147.0 | 96.47 |
| 14 | 18.19 | 10279 | 2237 | 22.6 | 55.10 | 14.80 | 4644 | 6635 | 41.9 | 96.29 |
| 15 | 22.47 | 14432 | 1804 | 14.2 | 60.83 | 18.47 | 10816 | 3085 | 38.4 | 94.15 |
| 16 | 38.52 | 20163 | 2471 | 25.1 | 60.28 | 27.71 | 8949 | 6448 | 60.8 | 94.67 |
| 17 | 40.72 | 17740 | 3683 | 40.5 | 58.43 | 29.54 | 8651 | 9334 | 67.5 | 95.64 |
| 18 | 57.06 | 23828 | 3901 | 41.7 | 64.69 | 39.50 | 9940 | 10143 | 76.4 | 95.29 |
| 19 | 35.45 | 16004 | 3472 | 43.7 | 54.52 | 25.99 | 7608 | 9149 | 66.2 | 95.59 |
| 20 | 43.45 | 18217 | 3945 | 39.6 | 59.13 | 31.28 | 8841 | 9276 | 63.8 | 95.68 |
| 21 | 28.37 | 13758 | 2951 | 23.8 | 52.03 | 21.00 | 5886 | 8353 | 81.1 | 95.62 |
| 22 | 69.45 | 30847 | 3437 | 41.6 | 57.94 | 50.24 | 11265 | 11295 | 71.6 | 95.36 |
| 23 | 36.76 | 17998 | 2806 | 29.7 | 46.29 | 28.75 | 7364 | 9287 | 81.5 | 96.20 |
| 24 | 49.34 | 22402 | 3254 | 30.2 | 64.52 | 34.97 | 9870 | 8227 | 66.4 | 95.41 |

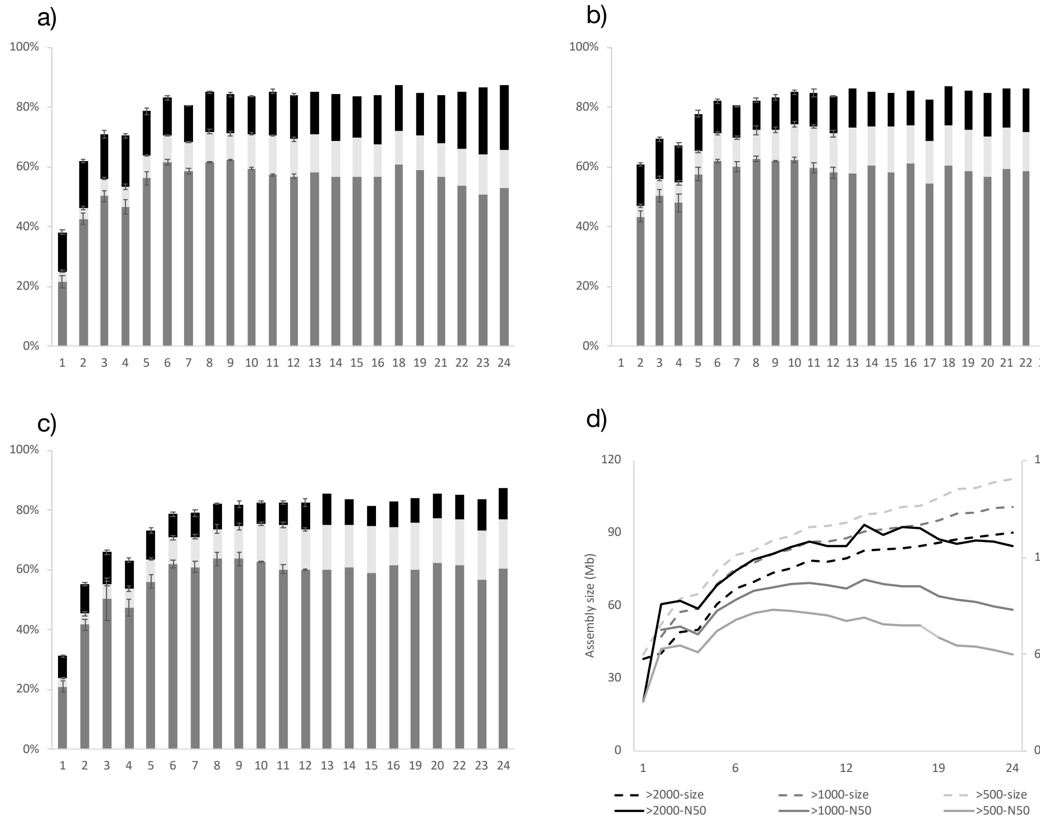

Figure S6. Summary statistics for different number of assembled nuclei (1-24) using assembly workflows 1 based on raw reads of individual nuclei assembled using Masurca, consensus assembly using Lingon. BUSCO estimates of completeness for a) workflow 1 contigs >500bp, b) workflow 1 contigs >1000bp, and c) workflow 1 contigs >2000bp. The later is the best option of the three, presented as method 1 in the main text. Percentage of single copy core genes detected as single copy (S: grey), duplicated (D: light grey) or fragmented (F: black). Average of 3-6 replicate assemblies up to 12 nuclei with error bars indicating SEM. In d) assembly size (dashed lines) and N50 (solid lines) for the there methods 1 >2000 bp (black), >10000 bp (grey) and >500 bp (light grey).

Table S6. Presence of the single copy genes EF1 and RPB1 in the generated assemblies. Present as single copy in both assembly methods 1 and 3, but not in assembly method 2. Contigs and regions where the gene is found are shown on the table.

| | EF1 | | RPB1 | |
|---|---|---|---|---|
| Assembly | Contig | Region | Contig | Region |
| 1 | contig006909 | 1349-2165 | contig004936 | 7862-10400 |
| 1n | contig003719 | 30811-31627 | contig003448 | 7862-10400 |
| 2 | contig010744 | 1152-1968,<br>14549-15365 | contig003808<br>contig001875<br>contig011350 | 7613-10151<br>1-1098<br>1-638 |
| 2n | scaffold_376 | 49952-50763,<br>63349-64165 | scaffold_1261<br>scaffold_339<br>scaffold_1645 | 9366-11904<br>36857-37954<br>3258-3895 |
| 3 | contig000025 | 24541-25357 | contig000552 | 15574-18112 |
| 3n | scaffold_129 | 24541-25357 | scaffold_994 | 15574-18112 |
